# Lasagna: Multifaceted Protein-Protein Interaction Prediction Based on Siamese Residual RCNN

**DOI:** 10.1101/501791

**Authors:** Muhao Chen, Chelsea J.-T. Ju, Guangyu Zhou, Tianran Zhang, Xuelu Chen, Kai-Wei Chang, Carlo Zaniolo, Wei Wang

**Affiliations:** Department of Computer Science, University of California, Los Angeles, 90095, USA; Department of Bioengineering, University of California, Los Angeles, 90095, USA

**Keywords:** PPI, deep learning, RCNN, neural sequence pair modeling

## Abstract

Sequence-based protein-protein interaction (PPI) prediction represents a fundamental computational biology problem. To address this problem, extensive research efforts have been made to extract predefined features from the sequences. Based on these features, statistical algorithms are learned to classify the PPIs. However, such explicit features are usually costly to extract, and typically have limited coverage on the PPI information. Hence, we present an end-to-end framework, Lasagna, for PPI predictions using only the primary sequences of a protein pair. Lasagna incorporates a deep residual recurrent convolutional neural network in the Siamese learning architecture, which leverages both robust local features and contextualized information that are significant for capturing the mutual influence of protein sequences. Our framework relieves the data pre-processing efforts that are required by other systems, and generalizes well to different application scenarios. Experimental evaluations show that Lasagna outperforms various state-of-the-art systems on the binary PPI prediction problem. Moreover, it shows a promising performance on more challenging problems of interaction type prediction and binding affinity estimation, where existing approaches fall short. The implementation of our framework is available at https://github.com/muhaochen/seq_ppi.git

## 1 Introduction

Detecting protein-protein interactions (PPIs) and characterizing the interaction types are essential toward understanding cellular biological processes in normal and disease states. Knowledge from these studies potentially facilitates therapeutic target identification [37] and novel drug design [45]. High-throughput experimental technologies have been rapidly developed to discover and validate PPIs in a large scale. These technologies include yeast two-hybrid screens [11], tandem affinity purification [12], and mass spectrometric protein complex identification [17]. However, experiment-based methods remain expensive, labor-intensive, and time-consuming. Most importantly, they often suffer from high levels of false-positive predictions [48, 56]. Evidently, there is an immense need for reliable computational approaches to identify and characterize PPIs.

The amino acid sequence represents the primary structure of a protein, which is the simplest type of information either obtained through direct sequencing or translated from DNA sequences. Many research efforts address the PPI problem based on predefined features extracted from protein sequences, such as ontological features of animo acids [20], autocovariance (AC) [13], conjoint triads (CT) [43] and composition-transition-distribution (CTD) descriptors [53]. These features generally summarize specific aspects of protein sequences such as physicochemical properties, frequencies of local patterns, and the positional distribution of amino acids. On top of these features, several statistical learning algorithms [13, 19, 30, 56, 58, 61] are applied to predict PPIs in the form of binary classification. These approaches provide feasible solutions to the problem. However, the extracted feature sets used in these approaches only have limited coverage on interaction information, as they are dedicated to specific facets of the protein profiles.

To mitigate the inadequacy of statistical learning methods, deep learning algorithms provide the powerful functionality to process large-scale data and automatically extract useful features for objective tasks [24]. Recently, deep learning architectures have produced powerful systems to address several bioinformatics problems related to single nucleotide sequences, such as genetic variants detection [2], DNA function classification [38], RNA-binding site prediction [60] and chromatin accessibility prediction [28]. These works typically use convolutional neural networks (CNN) [2, 60] for automatically selecting local features, or recurrent neural networks (RNN) [38] that aim at preserving the contextualized and long-term ordering information. By contrast, fewer efforts (discussed in Related Work) have been made to capture the pairwise interactions of proteins with deep learning, which remains a non-trivial problem with the following challenges: (i) Characterization of the proteins requires a model to effectively filter and aggregate their local features, while preserving significant contextualized and sequential information of the amino acids; (ii) Extending a deep neural architecture often leads to inefficient learning processes, and suffers from the notorious vanishing gradient problem [36]; (iii) An effective mechanism is also needed to apprehend the mutual influence of protein pairs in PPI prediction. Moreover, it is essential for the framework to be scalable to large data, and provides the potential to be generalized to different prediction tasks.

In this paper, we introduce Lasagna, a deep learning framework for PPI prediction using only the primary sequences of a protein pair. Lasagna employs a Siamese architecture to capture the mutual influence of a protein sequence pair. The learning architecture is based on a recurrent convolutional neural network (RCNN), which integrates multiple occurrences of convolution layers and residual gated recurrent units. To represent each amino acid in this architecture, Lasagna applies an efficient property-aware lexicon embedding approach to better capture the contextual and physicochemical relatedness of amino acids. This comprehensive encoding architecture provides a multi-granular feature aggregation process to effectively leverage both sequential and robust local information of the protein sequences. It is important to note that the scope of this work focuses only on the primary sequence as it is the fundamental information to describe a protein.

Our contributions are three-fold. First, we construct an end-to-end framework for PPI prediction that relieves the data pre-processing efforts for users. Lasagna requires only the primary protein sequences as the input, and is trained to automatically preserve the critical features from the sequences. Second, we emphasize and demonstrate the needs of considering the contextualized and sequential information when modeling the PPIs. Third, the architecture of Lasagna can be flexibly used to address different PPI tasks. Besides the binary prediction that is widely attempted in previous works, our framework extends its use to two additional challenging problems: multi-class interaction type prediction and binding affinity estimation. We use five datasets to evaluate the performance of our framework on these tasks. Lasagna outperforms various state-of-the-art approaches on the binary prediction task, which confirms the effectiveness in terms of integrating both local features and sequential information. The promising performance of the other two tasks demonstrates a wide usability of our approach. Especially on the binding affinity estimation of mutated proteins, Lasagna is able to respond to the subtle changes of point mutations and provides the best estimation with the smallest errors.

## 2 Related Work

Sequence-based approaches provide a critical solution to the binary PPI prediction task. Such works address the task with statistical learning models, including SVM [13, 58], kNN [53], Random Forest, multi-layer perceptron (MLP) [10] and ensemble ELM (EELM) [57]. These approaches rely on several feature extraction processes for the protein sequences, such as CT [48, 57], AC [13, 48, 57], CTD [10, 53], multi-scale continuous and discontinuous (MCD) descriptors [57], and local phase quantization (LPQ) [52]. These features measure physicochemical properties of the 20 canonical amino acids, and aim at summarizing full sequence information relevant to PPIs. More recent works [48, 51] propose the use of stacked autoencoders (SAE) to refine these heterogeneous features in lowdimensional spaces, which improve the aforementioned models on the binary prediction task. On the contrary, fewer efforts have been made towards multi-class prediction to infer the interaction types [44, 64] and the regression task to estimate binding affinity [47, 59]. These methods have largely relied on their capability of extracting and selecting better features, while the extracted features are far from fully exploiting the interaction information.

By nature, the PPI prediction task is comparable to the neural sentence pair modeling tasks in natural language processing (NLP) research, as they both seek to characterize the mutual influence of two sequences based on their latent features. In NLP, neural sentence pair models typically focus on capturing the discourse relations of lexicon sequences, such as textual entailment [18, 42, 55], paraphrases [15, 54] and sub-topic relations [5]. Many recent efforts adopt a Siamese encoding architecture, where encoders based on convolutional neural networks (CNN) [18, 54] and recurrent neural networks (RNN) [42] are widely used. A binary downstream classifier is then stacked to the sequence pair encoder for the detection of a targeted discourse relation. In contrast to sentences, proteins are profiled in sequences with more intractable patterns, as well as in a drastically larger range of lengths. Precisely capturing the PPI requires much more comprehensive learning architectures to refine the robust information from the entire sequences, and to preserve the long-term ordering information. One recent work [14], DPPI, uses a deep CNN-based architecture which focuses on capturing local features from protein profiles. DPPI represents the first work to deploy deep-learning to PPI prediction, and has achieved the state-of-the-art performance on the binary prediction task. However, it requires excessive efforts for data pre-processing such as constructing protein profiles by PSI-BLAST [1], and does not incorporate a neural learning architecture that captures the important contextualized and sequential features.

## 3 Methods

In this section, we introduce an end-to-end framework for sequence-based PPI prediction tasks. We begin with the denotations and problem specifications.

### 3.1 Preliminary

We use *A* to denote the vocabulary of 20 canonical amino acids. A protein is profiled as a sequence of amino acids *S* = [*a*_1_, *a*_2_, …, *a_l_*] such that each *a_i_* ∈ *A*. For each amino acid *a_i_*, we use bold-faced *a_i_* to denote its embedding representation, which we are going to specify in Section 3.2. We use *I* to denote the set of protein pairs, and *p* = (*S*_1_, *S*_2_) ∈ *I* denotes a pair of proteins of which our framework captures the interaction information.

We address three challenging PPI prediction tasks based only on the primary sequence information: (i) *Binary prediction* seeks to provide a binary classifier to indicate whether the corresponding protein pair interacts, which is the simplest and widely considered problem setting in previous works [14, 45, 48]. (ii) *Interaction type prediction* is a multi-class classification problem, which seeks to identify the interaction type of two proteins. (iii) *Binding affinity estimation* aims at producing a regression model to estimate the strength of the binding interaction.

### 3.2 RCNN-based Protein Sequence Encoder

We employ a deep Siamese architecture of Residual RCNN to capture latent semantic features of the protein sequence pairs.

#### Residual RCNN

The RCNN seeks to leverage both the global sequential information and local features that are significant to the characterization of PPI from the protein sequences. This deep neural encoder stacks multiple instances of two computational modules, i.e. *convolution layers with pooling* and *bidirectional residual gated recurrent units.*

#### The convolution layer with pooling

We use *X* = [v_1_, v_2_,…, v*_l_*] to denote an input vector sequence that corresponds to an embedded protein sequence or the outputs of a previous neural layer. A convolution layer applies a weight-sharing kernel 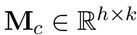 to generate a latent representation 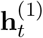 from a window **v**_*t:t*+*h*-1_ of the input vector sequence *X*:

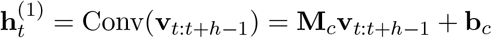

for which *h* is the kernel size and **b***_c_* is a bias vector. The convolution layer applies the kernel as a sliding window to produce a sequence of latent vectors 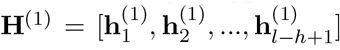, where each latent vector combines the local features from each *h*-gram of the input sequence. The *n*-max-pooling mechanism is applied to every consecutive *n*-length subsequence (i.e., non-overlapped *n*-stride) of the convolution outputs by 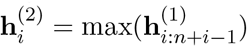. The purpose of this mechanism is to discretize the convolution results, and preserve the most significant features within each n-stride [5, 14, 23]. By definition, this mechanism divides the size of processed features by *n*.

#### The Residual Gated Recurrent Units

The Gated Recurrent Unit model (GRU) represents an alternative to the Long-short-term Memory network (LSTM) [6], which consecutively characterizes the sequential information without using separated memory cells [9]. Each unit consists of two types of gates to track the state of the sequence, i.e. the reset gate **r***_t_* and the update gate **z***_t_*. Given the vector representation **v***_t_* of an incoming item (either a pre-trained amino acid embedding, or an output of the previous layer), GRU updates the hidden state 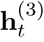 of the sequence as a linear combination of the previous state 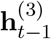 and the candidate state 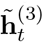 of a new item **v**_t_, which is calculated as below.

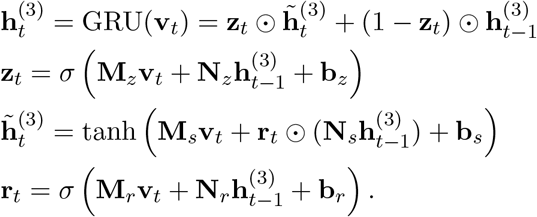

⨀ thereof denotes the element-wise multiplication. The update gate **z***_t_* balances the information of the previous sequence and the new item, where capitalized **M**_*_ and **N**_*_ denote different weight matrices, **b**_*_ denote bias vectors, and σ is the sigmoid function. The candidate state 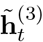 is calculated similarly to those in a traditional recurrent unit, and the reset gate **r***_t_* controls how much information of the past sequence contributes to 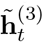. Note that GRU generally performs comparably to LSTM in sequence encoding tasks, but is less complex and requires much fewer computational resources for training [7].

A *bidirectional GRU layer* characterizes the sequential information in two directions. It contains the forward encoding process 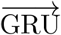 that reads the input vector sequence *X* = [**v**_1_, **v**_2_,…, **v**_*l*_] from **v**_l_ to **v***_l_*, and a backward encoding process 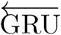 that reads in the opposite direction. The encoding results of both processes are concatenated for each input item 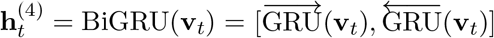.

The *residual mechanism* passes on an identity mapping of the GRU inputs to its output side through a residual shortcut [16]. By adding the forwarded input values to the outputs, the corresponding neural layer is only required to capture the difference between the input and output values. This mechanism aims at improving the learning process of non-linear neural layers by increasing the sensitivity of the optimization gradients [16, 22], as well as preventing the model from the vanishing gradient problem. It has been widely deployed in deep learning architectures for various tasks of image recognition [16], document classification [8] and speech recognition [62]. In our deep RCNN, the bidirectional GRU is incorporated with the residual mechanism, and will pass on the following outputs to its subsequent neural network layer:

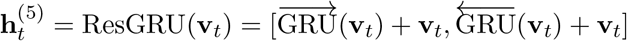

In our development, we have found that the residual mechanism is able to drastically simplify the training process, and largely decreases the epochs of parameter updates for the model to converge.

#### Protein sequence encoding

The RCNN encoder *E_RCNN_*(*S*) alternately stacks multiple occurrences of the above two intermediary neural network components. A convolution layer serves as the first encoding layer to extract local features from the input sequence. On top of that, a residual GRU layer takes in the preserved local features, whose outputs are passed to another convolution layer. Repeating of these two components in the network structure conducts an automatic multi-granular feature aggregation process on the protein sequence, while preserving the sequential and contextualized information on each granularity of the selected features. The last residual GRU layer is followed by another convolution layer for a final round of local feature selection to produce the last hidden states 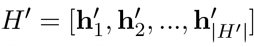. Note that the dimensionality of the last hidden states does not need to equal that of the previous hidden states. A high-level sequence embedding of the entire protein sequence is obtained from the global average-pooling [25] of 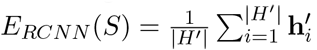. Fig. 1 shows the entire structure of our RCNN encoder.

**Fig. 1:**
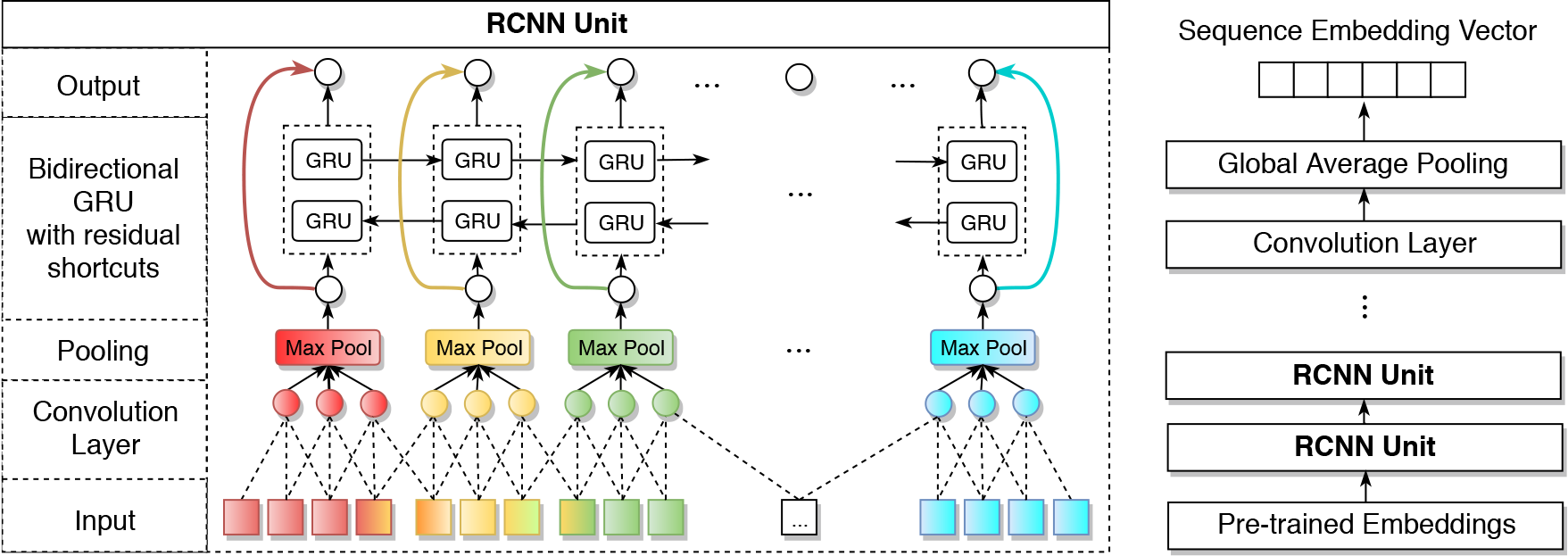
The structure of our residual RCNN encoder is shown on the right, and the RCNN unit is shown on the left. Each RCNN unit contains a convolution-pooling layer followed a bidirectional residual GRU.

#### Pre-trained Amino Acid Embeddings

To support inputting the non-numerical sequence information, we provide a useful embedding method to represent each animo acid *a* ∈ *A* as a semi-latent vector **a**. Each embedding vector is a concatenation of two sub-embeddings, i.e. **a** = [**a***_c_*, **a***_ph_*]. The first part **a***_c_* measures the co-occurrence similarity of the amino acids in protein sequences, which is obtained by pre-training the Skip-Gram model [27] on a collection of protein sequences. The second part **a***_ph_* represents the similarity of electrostaticity and hydrophobicity among amino acids. The 20 amino acids can be clustered into seven classes based on their dipoles and volumes of the side chains to reflect this property. Thus, **a***_ph_* is a one-hot encoding based on the classification defined by Shen et al. [43].

### 3.3 Learning Architecture and Learning Objectives

Our framework characterizes the interactions of protein pairs in the following two stages.

#### Siamese Architecture

The overall architecture of Lasagna is shown in Fig. 2. Given a pair of proteins *p* = (*S*_1_, *S*_2_) ∈ *I*, the same RCNN encoder is used to obtain the sequence embeddings *E_RCNN_*(*S*_1_) and *E_RCNN_*(*S*_2_) of both proteins. Both sequence embeddings are combined using element-wise multiplication, i.e., *E_RCNN_*(*S*_1_) ⨀ *E_RCNN_*(*S*_2_). This is a commonly used operation to infer the relation of sequence embeddings [14, 21, 40, 50]. Note that some works use the concatenation of sequence embeddings [48, 54] instead of their multiplication, which we find to be less effective in modeling the symmetric relations of proteins.

**Fig. 2:**
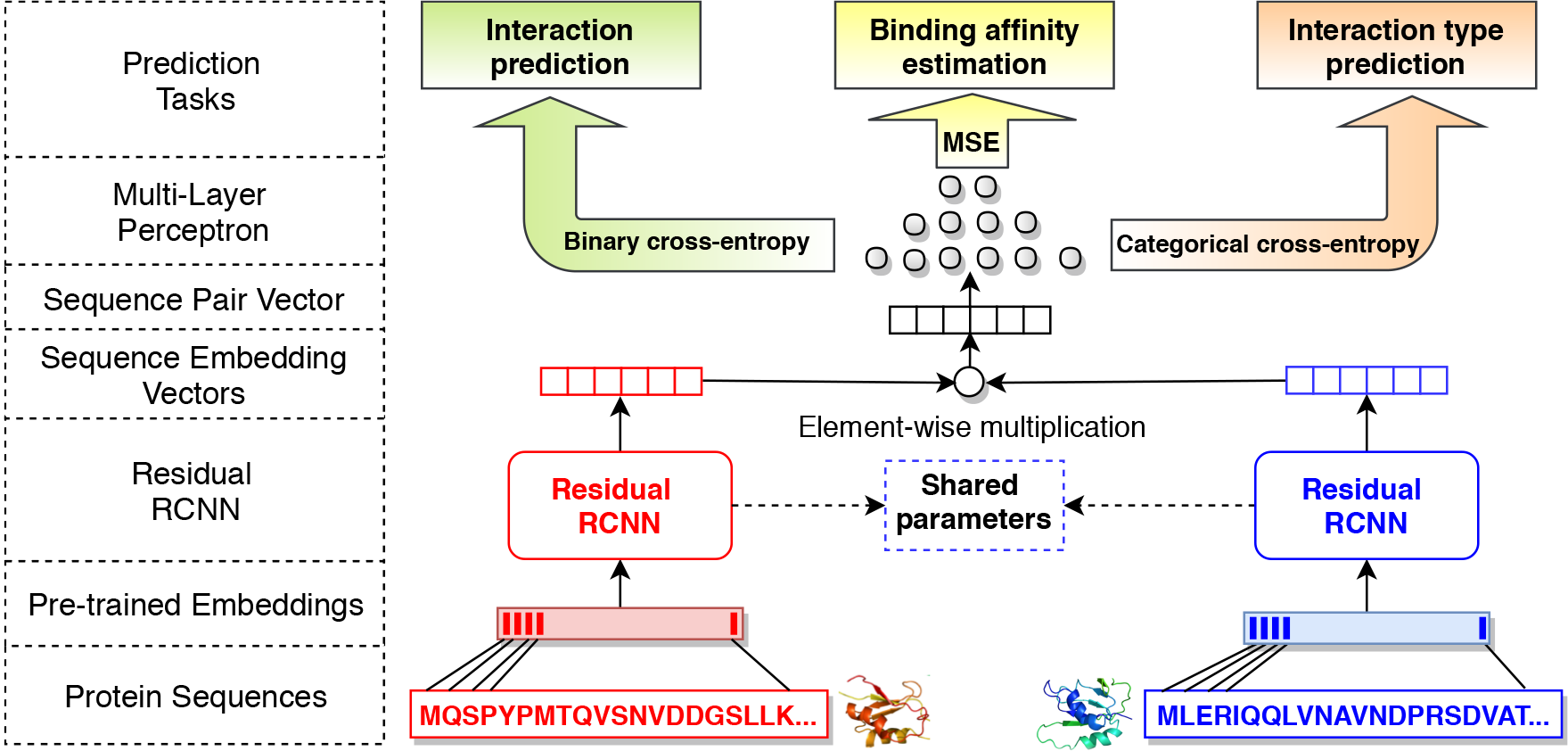
The overall learning architecture of our framework.

#### Learning objectives

A multi-layer perceptron (MLP) with leaky ReLU [26] is applied to the previous sequence pair representation, whose output 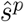 is either a vector or a scalar, depending on whether the model solves a classification or a regression task for the protein pair *p*. The entire learning architecture is trained to optimize the following two types of losses according to different PPI prediction problems.

(*i*) *Cross-entropy loss* is optimized for two classification problems, i.e. binary prediction and multi-class interaction type prediction. In this case, the MLP output 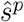 is a vector, whose dimensionality equals the number of classes 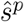 is normalized by a softmax function, where the *i*-th dimension 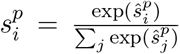 corresponds to the confidence score for the *i*-th class. The learning objective is to minimize the following cross-entropy loss, where *c^p^* is a one-hot indicator for the class label of protein pair *p*.

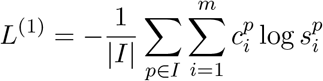

(*ii*) *Mean squared loss* is optimized for the binding affinity estimation task. In this case, 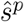 is a scalar output that is normalized by a sigmoid function 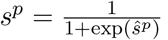, which is trained to approach the normalized ground truth score *c^p^* ∈ [0,1] by minimizing the following objective function:

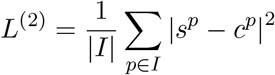

## 4 Experiments

We present the experimental evaluation of the proposed framework on three PPI prediction tasks, i.e. binary prediction, multi-class interaction type prediction, and binding affinity estimation.

### 4.1 Datasets

The experiments are conducted on the following datasets.

#### Guo’s Yeast PPI Dataset

Guo et al. [13] generate this dataset for the binary prediction of PPIs. There are 2,497 proteins forming 11,188 cases of PPIs, with half of them representing the positive samples, and the other half the negative samples. The positive samples are retrieved from the database of interacting proteins (DIP) [41]. The negative samples are generated by randomly pairing the proteins without evidence of interaction, and filtered by their subcellular locations. In other words, non-interactive pairs residing in the same location are excluded. The protein pairs and sequence information are available through the online supplementary materials^1^ of You et al. [56].

#### Pan’s Human PPI Dataset

Pan et al. [35] provide a set of data^2^ with 8,958 proteins representing both positive and negative interactions. The positive samples contain 36,630 interaction pairs retrieved from the human protein reference database (2007 version of HPRB [29]). Based on the assumption that two proteins are less likely to interact if they reside far away from each other, the negative samples are generated by pairing those found in different subcellular locations. Besides this pairing process, additional negative samples of human PPIs are included from the Negatome database [4], resulting in a total of 36,480 non-interaction pairs. We also use this dataset to evaluate the binary prediction task for PPIs.

#### STRING Datasets

The STRING database [49] annotates PPIs with their types. There are seven types of interactions: activation, binding, catalysis, expression, inhibition, post-translational modification (ptmod), and reaction. We download all interaction pairs for *Homo sapiens* from database version 10.5 [49]. Among the corresponding proteins, we randomly select 3,000 proteins and 8,000 proteins to generate two subsets. In this process, we randomly sample instances of different interaction types to ensure a balanced class distribution. Eventually, the two generated datasets, denoted by SHS27k and SHS148k, contain 26,945 cases and 148,051 cases of interactions respectively. We use these two datasets for the evaluation of the PPI type prediction task.

#### SKEMPI Dataset

We obtain the protein binding affinity data from SKEMPI (the Structural database of Kinetics and Energetics of Mutant Protein Interactions) [31] for the affinity estimation task. It contains 3,047 binding affinity changes upon mutation of protein sub-units within a protein complex. The binding affinity is measured by equilibrium dissociation constant (*K_d_*), reflecting the strength of biomolecular interactions. The smaller *K_d_* value means the higher binding affinity. Each protein complex contains single or multiple amino acid substitutions. The sequence of the protein complex is retrieved from the Protein Data Bank (PDB) [3]. We manually replace the mutated amino acids. For duplicate entries, we take the average *K_d_*. The final dataset results in the binding affinity of 2,792 mutant protein complexes, along with 158 wild-type.

### 4.2 Binary PPI Prediction

Binary PPI prediction is the primary task targeted by a handful of previous works [14, 43, 48, 53, 56]. The objective of these works is to identify whether a given pair of proteins interacts or not based on their primary sequences. We evaluate Lasagna based on the Yeast and the Human datasets.

Our model is compared against multiple baseline approaches, including: SVM-AC [13], kNN-CTD [53], EELM-PCA [57], SVM-MCD [58], MLP [10], Random Forest LPQ (RF-LPQ) [52] and DPPI [14]. Meanwhile, we also report the results of a Siamese Residual GRU (SRGRU) architecture, which is a simplification of Lasagna, where we discard all intermediary convolution layers and keep only the bidirectional residual GRU. The purpose of SRGRU is to show the significance of the contextualized and sequential information of protein profiles in characterizing PPIs. We also report the results of Siamese CNN (SCNN) by removing the residual GRU in Lasagna. This degenerates our framework to a similar architecture to DPPI, but differs in directly conducting an end-to-end training on primary sequences instead of requiring the protein profiles constructed by PSI-BLAST.

We use AMSGrad [39] to optimize the cross-entropy loss, for which we set the learning rate *α* to 0.001, the exponential decay rates *β*_1_ and *β*_2_ to 0.9 and 0.999, and batch size to 256 on both datasets. The number of occurrences for the RCNN units (i.e., one convolution-pooling layer followed by one bidirectional residual GRU layer) is set to 5, where we adopt 3-max-pooling and the convolution kernel of size 3. We set the hidden state size to be 50, and the RCNN output size to be 100. We set this configuration to ensure the RCNN to compress the selected features in a reasonably small vector sequence, before the features are aggregated by the last global average-pooling. Note that the model configuration study is provided in Appendix I. All model variants are trained until converge at each fold of the cross-validation. For amino acid embeddings, we pre-train the Skip-Gram on all sequences on our largest STRING dataset, SHS148k, using a context window size of 6 and a negative sampling size of 5. This process obtains an 5-dimensional vector for the first sub-embedding part of each amino acid, which is concatenated to the dipoles and side chain volume based one-hot encoding (**a***_ph_*) to obtain the 12-dimensional embedding representation of amino acids. We also evaluate different representations in Appendix II. As reported in Table A2 of Appendix III, the embedding pre-training process finishes within 1 minute on a commodity workstation, which is a one-time effort that can be reused on different tasks and datasets. The obtained amino acid embeddings are used to transform the protein sequences to vector sequences, for which we zero-pad short sequences to the highest sequence length in the dataset. This is a widely adopted technique for sequence modeling in NLP [5, 15, 18, 42, 55, 63] as well as in bioinformatics [28, 33, 34] for efficient training.

#### Evaluation protocol

Following the settings in previous works [14, 43, 48, 56, 58], we conduct 5-fold cross-validation on the Yeast dataset and 10-fold cross-validation on the Human dataset. We aggregate three metrics on the test cases of each fold, i.e. the overall *accuracy, precision* and *F1* on positive cases. All these metrics are preferred to be higher to indicate better performance.

#### Results

As shown in Fig. 3, the CNN-based architecture, DPPI, demonstrates state-of-the-art performance over other baselines that employ statistical learning algorithms or densely connected MLP. This shows the superiority of deep-learning-based techniques in encapsulating various types of information of a protein pair, such as amino acid composition and their co-occurrences, and automatically extracting the robust ones for the learning objectives. That said, DPPI requires an extensive effort in data pre-processing, specifically in constructing the protein profile for each sequence. On average, each PSI-BLAST search of a protein requires around one hour of computation on our server, and we are unable to obtain the profile for 8,958 protein sequences in the Human dataset to evaluate DPPI. For this reason, we implement SCNN to evaluate the performance of a simplified CNN architecture on different datasets. Our SCNN implementation produces comparable results as DPPI while evaluating the Yeast dataset. Compared to the use of SAE [48], Fig. 4 shows that the simplified version of our framework with only CNN can already leverage the significant features from primary protein sequences. In addition, the SRGRU architecture has offered comparable performance to SCNN on both datasets. This indicates that preserving the sequential and contextualized features of the protein sequences is as crucial as incorporating the local features. By integrating both significant local features and sequential information, Lasagna outperforms DPPI by 2.68% in accuracy, 0.61% in precision, and 2.83% in F1 on the Yeast dataset, and outperforms SCNN by 0.90% in accuracy, 1.78% in precision, and 1.01% in F1 on the Human dataset. Thus, the residual RCNN is very promising for modeling binary PPIs.

**Fig. 3:**
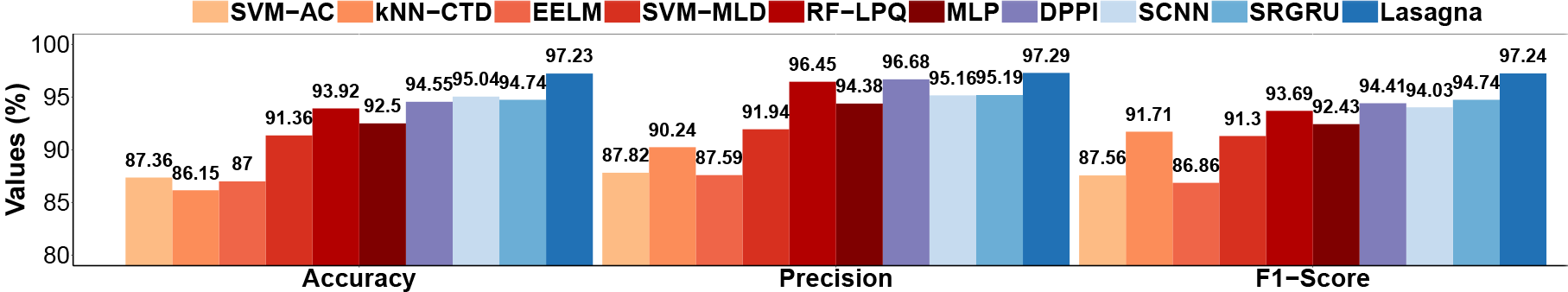
Evaluation of binary PPI prediction on the Yeast dataset.

**Fig. 4:**
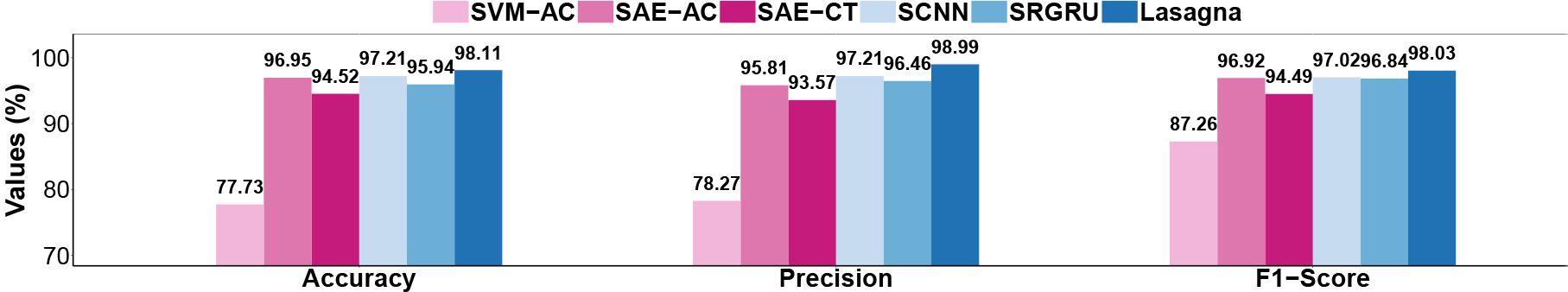
Evaluation of binary PPI prediction on the Human dataset.

### 4.3 Interaction Type Prediction

The objective of this task is to predict the interaction type of two interacting proteins. We evaluate this task based on SHS27k and SHS148k datasets. To the best of our knowledge, much fewer efforts attempt for the multi-class PPI prediction in contrast to the binary prediction. Zhu et al. [64] train a two-stage SVM classifier to distinguish obligate, non-obligate, and crystal packing interactions; Silberberg et al. [44] use logistic regression to predict several types of enzymetic actions. However, none of their implementations are publicly available. Different from the categories of interaction types used above, we aim at predicting the interaction types annotated by the STRING database.

Table 1: Accuracy (%) and fold changes over zero rule for PPI interaction type prediction on two STRING datasets.

**Table 1:**
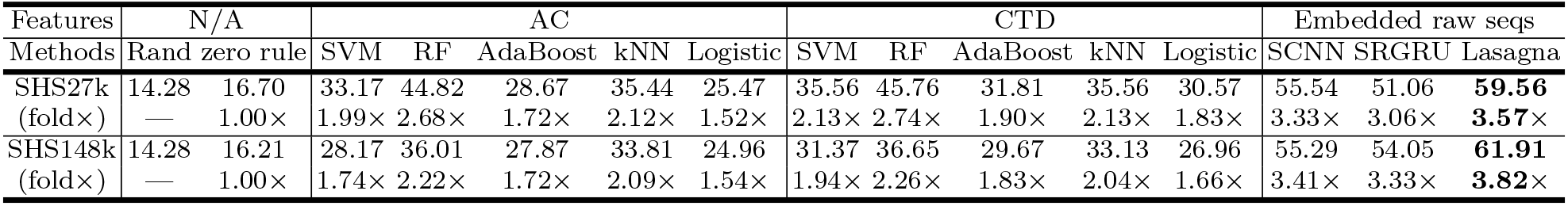
Accuracy (%) and fold changes over zero rule for PPI interaction type prediction on two STRING datasets.

**Table 2:** Results for binding affinity prediction on the SKEMPI dataset.

| Features | AC |  |  |  | CTD |  |  |  | Embedded raw seqs |  |  |
| --- | --- | --- | --- | --- | --- | --- | --- | --- | --- | --- | --- |
| Methods | BR | SVM | RF | AdaBoost | BR | SVM | RF | AdaBoost | SCNN | SRGRU | Lasagna |
| $MSE(\times 10^{-2})$ | 1.70 | 2.20 | 1.77 | 1.98 | 1.86 | 1.84 | 1.49 | 1.84 | 0.87 | 0.95 | **0.63** |
| $MAE(\times 10^{-2})$ | 9.56 | 11.81 | 9.81 | 11.15 | 10.20 | 11.04 | 9.06 | 10.69 | 6.49 | 7.08 | **5.48** |
| $Corr$ | 0.564 | 0.353 | 0.546 | 0.451 | 0.501 | 0.501 | 0.640 | 0.508 | 0.831 | 0.812 | **0.873** |

We train several statistical learning algorithms on the widely employed AC and CTD features for protein characterization as our baselines. These algorithms include SVM, Random Forest, Adaboost (SAMME.R algorithm [65]), the kNN classifier and logistic regression. For deep-learning-based approaches, we deploy the SCNN architecture where an output MLP with categorical cross-entropy loss is incorporated, as well as a similar SRGRU architecture into comparison. Results of two naive baselines of random guessing and zero rule (i.e., simply predicting the majority class) are also reported for reference.

#### Evaluation protocol

All approaches are evaluated on the two datasets by 10-fold cross-validation, using the same partition scheme for an unbiased evaluation. We carry forward the model configurations from the last experiment to evaluate the performance of the frameworks under controlled variables. For baseline models, we examine three different ways of combining the feature vectors of the two input proteins, i.e. element-wise multiplication, the Manhattan difference (i.e. the absolute differences of corresponding features [32]) and concatenation. The Manhattan difference thereof, consistently obtains better performance, considering the small values of the input features and the asymmetry of the captured protein relations.

#### Results

The prediction accuracy and fold changes over the zero rule baseline are reported in Table 1. Note that since the multi-class prediction task is much more challenging than the binary prediction task, it is expected to observe lower accuracy than reported in the previous experiment. Among all the baselines using explicit features, the CTD-based models generally perform better than the AC-based ones. CTD descriptors seek to cover both of the continuous and discontinuous interaction information [53], which potentially better discriminate among PPI types.

The best baseline using Random Forest thereof, achieves satisfactory results by more than doubling the accuracy of zero rule on the smaller SHS27k dataset. However, on the larger SHS148k dataset, the accuracy of these explicit-feature-based models is notably impaired. We hypothesize that, such predefined explicit features are not representative enough to distinguish the PPI types. On the other hand, the deep-learning-based approaches do not need to explicitly utilize these features, and perform consistently well on both settings. The raw sequence information is sufficient for these approaches to drastically outperform the Random Forest by at least 5.30% in accuracy on SHS27k and 17.40% in accuracy on SHS148k. SCNN thereof outperforms SRGRU by 4.48% and 1.24% in accuracy on SHS27k and SHS148k, respectively. This implies that the local interacting features are relatively more deterministic than contextualized and sequential features on this task. The results by the residual RCNN-based framework are very promising, as it outperforms SCNN by 4.02% and 6.62% in accuracy on SHS27k and SHS148k respectively. It also remarkably outperforms the best explicit-feature-based baselines on the two datasets by 13.80% and 25.26% in accuracy, and more than 3.5 of fold changes over the zero rule on both datasets.

### 4.4 Binding Affinity Estimation

Lastly, we evaluate Lasagna for binding affinity estimation using the SKEMPI dataset. We employ the mean squared loss variant of Lasagna to address this regression task. Since the lengths of protein sequences in SKEMPI are much shorter than those in the other datasets, we accordingly reduce the occurrences of RCNN units to 3, while other configurations remain unchanged. For baselines, we compare against several regression models based on the AC and CTD features, which include Bayesian Redge regressor (BR), SVM, Adaboost with decision tree regressors and Random Forest regressor. The corresponding features for two sequences are again combined via the Manhattan difference. We also modify SCNN and SRGRU to their mean squared loss variants, in which we reduce the layers in the same way of RCNN.

#### Evaluation protocol

We aggregate three metrics through 10-fold cross-validation, i.e. *mean squared error* (*MSE*), *mean absolute error* (*MAE*) and *Pearson’s correlation coefficient* (*Corr*). These are three commonly reported metrics for regression tasks, for which lower *MSE* and *MAE* as well as higher *Corr* indicate better performance. In the cross-validation process, we normalize the affinity values of the SKEMPI dataset to [0,1] via min-max rescaling^3^.

#### Results

Table 2 reports the results for this experiment. It is noteworthy that, one single change of amino acid can lead to a drastic effect on binding affinity. While such subtle changes are difficult to be reflected by the explicit features, the deep-learning-based methods can competently capture such changes from the raw sequences. Our RCNN-based framework again offers the best performance among the deep-learning-based approaches, and significantly outperforms the best baseline (CTD-based Random Forest) by offering a 0.233 increase in *Corr*, as well as remarkably lower *MSE* and *MAE*. While our experiment is conducted on a relatively small dataset, we seek to extend our Lasagna framework to a more generalized solution for binding affinity estimation, once a larger and more heterogeneous corpus is available.

## 5 Conclusion and Future Work

In this paper, we propose a novel end-to-end framework for PPI prediction based on the primary sequences. Our proposed framework, Lasagna, employs a residual RCNN, which provides an automatic multi-granular feature selection mechanism to capture both local significant features and sequential features from the primary protein sequences. By incorporating the RCNN in a Siamese-based learning architecture, the framework captures effectively the mutual influence of protein pairs, and generalizes well to address different PPI prediction tasks without the need of predefined features. Extensive experimental evaluations on five datasets show promising performance of our framework on three challenging PPI prediction tasks. This also leads to significant amelioration over various baselines. Different sizes of the datasets also show satisfying scalability of the framework. For future work, one important direction is to apply the Lasagna framework to other sequence-based inference tasks in bioinformatics, such as modeling RNA and protein interactions. We also seek to extend multi-task learning [46] into our framework, and jointly learn different objectives based on corpora with multifaceted interaction information.

## Footnotes

1 https://doi.org/10.1371/journal.pone.0125811.s002

2 http://www.csbio.sjtu.edu.cn/bioinf/LR_PPI/Data.htm

3 https://en.wikipedia.org/wiki/Feature_scaling

## Appendix

**Fig. A1:**
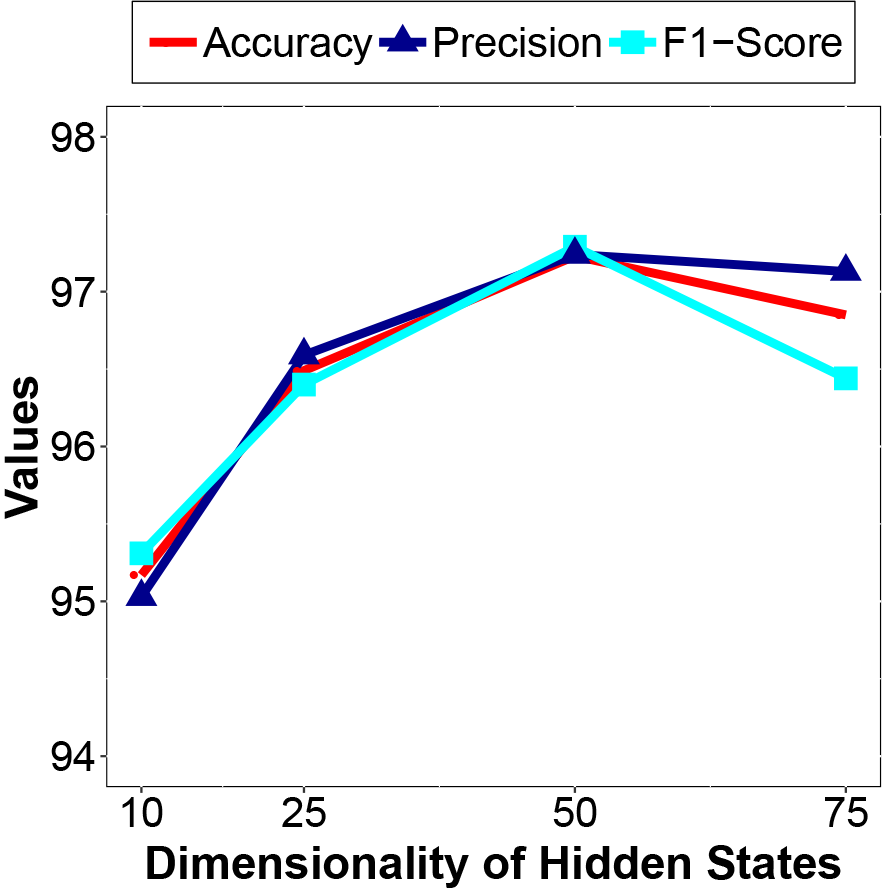
Performance evaluation on dimensionality of hidden states.

**Fig. A2:**
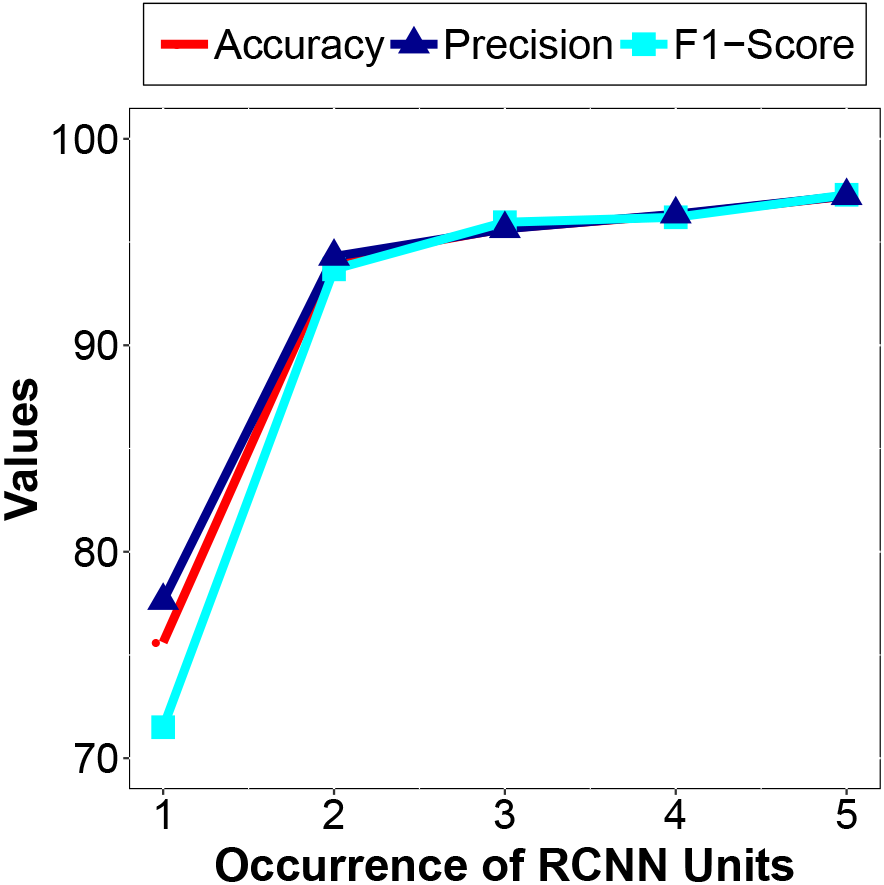
Performance evaluation on the number of occurrences of the RCNN units.

## I Model Configuration Study

We examine the configuration of two critical factors that can affect the performance of our framework: the dimensionality of hidden states and the number of occurrences for the RCNN units. We show the effects of different settings of these two factors based on the binary PPI prediction task on the Yeast dataset. The hidden state sizes are chosen from {10, 25, 50, 75}. As illustrated in Fig A1, the performance of Lasagna initially increases as we raise the dimensionality of the hidden states, but decreases when we set the dimensionality to 75. The occurrences of RCNN units contribute to the levels of granularity in feature aggregation. The fewer the occurrences corresponds to less aggregation. However, too many occurrences can lead to over-compressing the features. We examine the occurrences from 1 to 5 based on Yeast. Note that we do not adopt the setting with 6 occurrences, where the RCNN encoder over-compresses the extracted features to a very small number of latent vectors before the last global average pooling. Aligned with our hypothesis, Fig A2 shows that the accuracy, precision, and F1-score increase when we increase the number of occurrences of the RCNN units. The improvement from 2 to 5 occurrences is marginal, which shows that our framework is robust to this setting as long as there are more than 2 occurrences of RCNN units.

## II Comparison of Amino Acid Representations

We evaluate the effectiveness of different strategies in representing the 20 canonical amino acids in our framework using the Yeast dataset. As describe in Section 3.2, we use a pre-trained 12dimensional embedding to represent each amino acid. The first part of the embedding vector contains a 5-dimensional vector, **a***_c_*, which measures the co-occurrence similarity of the amino acids in protein sequences. The second part contains a 7-dimensional vector, **a***_ph_*, which describes the categorization of electrostaticity and hydrophobicity for the amino acid. We examine the performance of using each part individually, as well as the performance of combining them as used in our framework. In addition, we include a simple one-hot vector representation, which does not consider the relatedness of amino acids and treats each of them independently.

**Table A1:** Comparison of different amino acid representations.

| | $[a_c, a_{ph}]$ | $a_c$ only | $a_{ph}$ only | one-hot (20) |
| --- | --- | --- | --- | --- |
| **Accuracy** | **97.23** | 96.67 | 96.03 | 96.11 |
| **Precision** | **97.29** | 96.35 | 95.91 | 96.34 |
| **F1-Score** | **97.24** | 96.51 | 96.08 | 96.10 |

The results in Table A1 show that, once we remove either of the two parts of the proposed embedding, the performance of the model slightly drops. Meanwhile, the proposed pre-trained embeddings lead to noticeably better performance of the model than adopting the simple one-hot encodings of the canonical amino acids.

## III Run-time Analysis

We report training time for each experiment in Table A2, as well as the training time of the amino acid embeddings. All these processes are conducted on one NVIDIA GeForce GTX 1080 Ti GPU. The amino acid embeddings are trained on the 8,000 sequences of the SHS148k dataset. The training setup has been described in Section 4.2. The process is very efficient, which only takes 8 seconds to complete. This is also a one-time effort, as the same embeddings can be applied to different tasks and datasets.

For each experiment, we calculate the average training time over either 5-fold (Yeast dataset) or 10-fold (others) cross-validation. In both binary and multi-class predictions, the training time increases along with the increased number of training cases. The regression estimation generally requires more iterations per training case to converge than classification tasks. Thus, with much fewer cases, the training time on SKEMPI for affinity estimation is more than that on the Yeast dataset for binary prediction.

**Table A2:** Run-Time of embedding training and different prediction tasks.

| Task | Embedding Training | Binary Prediction | Binary Prediction | Multi-class Prediction | Multi-class Prediction | Regression |
| --- | --- | --- | --- | --- | --- | --- |
| **Dataset** | **SHS148k** | **Yeast** | **Human** | **SHS27k** | **SHS148k** | **SKEMPI** |
| **Sample Size** | 8,000 | 11,188 | 73,110 | 26,945 | 148,051 | 2,950 |
| **Run-Time** | 8sec | 2.5min | 19.2min | 15.8min | 138.3min | 12.5min |

## Bibliography

[1] S. F. Altschul, T. L. Madden, A. A. Schäffer, J. Zhang, Z. Zhang, W. Miller, and D. J. Lipman. Gapped blast and psi-blast: a new generation of protein database search programs. Nucleic acids research, 25(17):3389–3402, 1997.

[2] C. Anderson. Google’s ai tool deepvariant promises significantly fewer genome errors. Clinical OMICs, 5(1):33–33, 2018.

[3] H. M. Berman, J. Westbrook, Z. Feng, G. Gilliland, T. N. Bhat, H. Weissig, I. N. Shindyalov, and P. E. Bourne. The protein data bank. Nucleic acids research, 28(1):235–242, 2000.

[4] P. Blohm, G. Frishman, P. Smialowski, F. Goebels, B. Wachinger, A. Ruepp, and D. Frishman. Negatome 2.0: a database of non-interacting proteins derived by literature mining, manual annotation and protein structure analysis. Nucleic acids research, 42(D1):D396–D400, 2013.

[5] M. Chen, C. Meng, G. Huang, and C. Zaniolo. Neural article pair modeling for wikipedia sub-article matching. In European Conference on Machine Learning (ECML), 2018.

[6] K. Cho, B. van Merrienboer, C. Gulcehre, D. Bahdanau, F. Bougares, H. Schwenk, and Y. Bengio. Learning phrase representations using rnn encoder-decoder for statistical machine translation. In Proceedings of the Conference on Empirical Methods in Natural Language Processing (EMNLP), pages 1724–1734, 2014.

[7] J. Chung, C. Gulcehre, K. Cho, et al. Empirical evaluation of gated recurrent neural networks on sequence modeling. In Deep Learing@NIPS, 2014.

[8] A. Conneau, H. Schwenk, L. Barrault, and Y. Lecun. Very deep convolutional networks for text classification. In European Chapter of the Association for Computational Linguistics, 2017.

[9] B. Dhingra, H. Liu, et al. Gated-attention readers for text comprehension. In Proceedings of the 54th Annual Meeting of the Association for Computational Linguistics, 2016.

[10] X. Du, S. Sun, C. Hu, Y. Yao, Y. Yan, and Y. Zhang. Deepppi: boosting prediction of protein–protein interactions with deep neural networks. Journal of chemical information and modeling, 57(6):1499–1510, 2017.

[11] S. Fields and O.-k. Song. A novel genetic system to detect protein–protein interactions. Nature, 340(6230):245, 1989.

[12] A.-C. Gavin, M. Bösche, R. Krause, P. Grandi, M. Marzioch, A. Bauer, J. Schultz, J. M. Rick, A.-M. Michon, C.-M. Cruciat, et al. Functional organization of the yeast proteome by systematic analysis of protein complexes. Nature, 415(6868):141, 2002.

[13] Y. Guo, L. Yu, Z. Wen, and M. Li. Using support vector machine combined with auto covariance to predict protein-protein interactions from protein sequences. Nucleic acids research, 36(9):3025–30, may 2008.

[14] S. Hashemifar, B. Neyshabur, A. A. Khan, and J. Xu. Predicting protein-protein interactions through sequence-based deep learning. Bioinformatics, 34(17):i802–i810, 2018.

[15] H. He, K. Gimpel, and J. Lin. Multi-perspective sentence similarity modeling with convolutional neural networks. In Proceedings of the Conference on Empirical Methods in Natural Language Processing (EMNLP), pages 1576–1586, 2015.

[16] K. He, X. Zhang, S. Ren, and J. Sun. Deep residual learning for image recognition. In Proceedings of the IEEE conference on computer vision and pattern recognition, pages 770–778, 2016.

[17] Y. Ho, A. Gruhler, A. Heilbut, G. D. Bader, L. Moore, S.-L. Adams, A. Millar, P. Taylor, K. Bennett, K. Boutilier, et al. Systematic identification of protein complexes in saccharomyces cerevisiae by mass spectrometry. Nature, 415(6868):180, 2002.

[18] B. Hu, Z. Lu, H. Li, and Q. Chen. Convolutional neural network architectures for matching natural language sentences. In NIPS, pages 2042–2050, 2014.

[19] Y.-A. Huang, Z.-H. You, X. Gao, L. Wong, and L. Wang. Using weighted sparse representation model combined with discrete cosine transformation to predict protein-protein interactions from protein sequence. BioMed research international, 2015, 2015.

[20] R. Jansen, H. Yu, D. Greenbaum, Y. Kluger, N. J. Krogan, S. Chung, A. Emili, M. Snyder, J. F. Greenblatt, and M. Gerstein. A Bayesian networks approach for predicting protein-protein interactions from genomic data. Science, 302(5644):449–53, oct 2003.

[21] J.-Y. Jiang, F. Chen, Y.-Y. Chen, and W. Wang. Learning to disentangle interleaved conversational threads with a siamese hierarchical network and similarity ranking. In NAACL, 2018.

[22] J. Kim, J. Kwon Lee, and K. Mu Lee. Accurate image super-resolution using very deep convolutional networks. In Proceedings of the IEEE conference on computer vision and pattern recognition, pages 1646–1654, 2016.

[23] Y. Kim. Convolutional neural networks for sentence classification. In Proceedings of the Conference on Empirical Methods in Natural Language Processing (EMNLP), 2014.

[24] Y. LeCun, Y. Bengio, and G. Hinton. Deep learning. nature, 521(7553):436, 2015.

[25] M. Lin, Q. Chen, and S. Yan. Network in network. In ICLR, 2013.

[26] A. L. Maas, A. Y. Hannun, and A. Y. Ng. Rectifier nonlinearities improve neural network acoustic models. In ICML, volume 30, page 3, 2013.

[27] T. Mikolov, I. Sutskever, K. Chen, G. S. Corrado, and J. Dean. Distributed representations of words and phrases and their compositionality. In C. J. C. Burges, L. Bottou, M. Welling, Z. Ghahramani, and K. Q. Weinberger, editors, NIPS, pages 3111–3119. 2013.

[28] X. Min, W. Zeng, N. Chen, T. Chen, and R. Jiang. Chromatin accessibility prediction via convolutional long short-term memory networks with k-mer embedding. Bioinformatics, 33(14):i92–i101, 2017.

[29] G. R. Mishra, M. Suresh, K. Kumaran, N. Kannabiran, S. Suresh, P. Bala, K. Shivakumar, N. Anuradha, R. Reddy, T. M. Raghavan, et al. Human protein reference database2006 update. Nucleic acids research, 34(suppl_1):D411–D414, 2006.

[30] M. Miwa, R. Sætre, Y. Miyao, and J. Tsujii. A rich feature vector for protein-protein interaction extraction from multiple corpora. In EMNLP, pages 121–130. ACL, 2009.

[31] I. H. Moal and J. Fernández-Recio. Skempi: a structural kinetic and energetic database of mutant protein interactions and its use in empirical models. Bioinformatics, 28(20):2600–2607, 2012.

[32] J. Mueller and A. Thyagarajan. Siamese recurrent architectures for learning sentence similarity. In AAAI, volume 16, pages 2786–2792, 2016.

[33] A. T. Muller, J. A. Hiss, and G. Schneider. Recurrent neural network model for constructive peptide design. Journal of chemical information and modeling, 58(2):472–479, 2018.

[34] X. Pan and H.-B. Shen. Predicting rna-protein binding sites and motifs through combining local and global deep convolutional neural networks. Bioinformatics, 2018.

[35] X.-Y. Pan, Y.-N. Zhang, and H.-B. Shen. Large-scale prediction of human protein-protein interactions from amino acid sequence based on latent topic features. Journal of Proteome Research, 9(10):4992–5001, 2010.

[36] R. Pascanu, T. Mikolov, and Y. Bengio. On the difficulty of training recurrent neural networks. In International Conference on Machine Learning, pages 1310–1318, 2013.

[37] I. Petta, S. Lievens, C. Libert, J. Tavernier, and K. De Bosscher. Modulation of protein–protein interactions for the development of novel therapeutics. Molecular Therapy, 24(4):707–718, 2016.

[38] D. Quang and X. Xie. Danq: a hybrid convolutional and recurrent deep neural network for quantifying the function of dna sequences. Nucleic acids research, 44(11):e107–e107, 2016.

[39] S. J. Reddi, S. Kale, and S. Kumar. On the convergence of adam and beyond. In ICLR, 2018.

[40] T. Rocktaschel, E. Grefenstette, K. M. Hermann, T. Kocisky, and P. Blunsom. Reasoning about entailment with neural attention. In International Conference on Learning Representations (ICLR), 2016.

[41] L. Salwinski, C. S. Miller, A. J. Smith, F. K. Pettit, J. U. Bowie, and D. Eisenberg. The database of interacting proteins: 2004 update. Nucleic acids research, 32(suppl_1):D449–D451, 2004.

[42] L. Sha, B. Chang, et al. Reading and thinking: Re-read lstm unit for textual entailment recognition. In Proceedings of the International Conference on Computational Linguistics, 2016.

[43] J. Shen, J. Zhang, X. Luo, W. Zhu, K. Yu, K. Chen, Y. Li, and H. Jiang. Predicting protein–protein interactions based only on sequences information. Proceedings of the National Academy of Sciences of the United States of America, 104(11):4337–41, mar 2007.

[44] Y. Silberberg, M. Kupiec, and R. Sharan. A method for predicting protein-protein interaction types. PLoS One, 9(3):e90904, 2014.

[45] L. Skrabanek, H. K. Saini, G. D. Bader, and A. J. Enright. Computational prediction of protein–protein interactions. Molecular biotechnology, 38(1):1–17, 2008.

[46] V. Smith, C.-K. Chiang, M. Sanjabi, and A. S. Talwalkar. Federated multi-task learning. In NIPS, pages 4424–4434, 2017.

[47] Y. S. Srinivasulu, J.-R. Wang, K.-T. Hsu, M.-J. Tsai, P. Charoenkwan, W.-L. Huang, H.-L. Huang, and S.-Y. Ho. Characterizing informative sequence descriptors and predicting binding affinities of heterodimeric protein complexes. BMC bioinformatics, 16(18):S14, 2015.

[48] T. Sun, B. Zhou, L. Lai, and J. Pei. Sequence-based prediction of protein protein interaction using a deep-learning algorithm. BMC bioinformatics, 18(1):277, 2017.

[49] D. Szklarczyk, J. H. Morris, H. Cook, M. Kuhn, S. Wyder, M. Simonovic, A. Santos, N. T. Doncheva, A. Roth, P. Bork, et al. The string database in 2017: quality-controlled protein–protein association networks, made broadly accessible. Nucleic acids research, page gkw937, 2016.

[50] K. S. Tai, R. Socher, and C. D. Manning. Improved semantic representations from tree-structured long short-term memory networks. In ACL, volume 1, pages 1556–1566, 2015.

[51] Y.-B. Wang, Z.-H. You, X. Li, T.-H. Jiang, X. Chen, X. Zhou, and L. Wang. Predicting protein–protein interactions from protein sequences by a stacked sparse autoencoder deep neural network. Molecular BioSystems, 13(7):1336–1344, 2017.

[52] L. Wong, Z.-H. You, S. Li, Y.-A. Huang, and G. Liu. Detection of protein-protein interactions from amino acid sequences using a rotation forest model with a novel pr-lpq descriptor. In ICIC, pages 713–720, 2015.

[53] L. Yang, J.-F. Xia, and J. Gui. Prediction of protein-protein interactions from protein sequence using local descriptors. Protein and Peptide Letters, 17(9):1085–1090, 2010.

[54] W. Yin and H. Schütze. Convolutional neural network for paraphrase identification. In Proceedings of the Conference of the North American Chapter of the Association for Computational Linguistics, pages 901–911, 2015.

[55] W. Yin, H. Schütze, et al. Abcnn: Attention-based convolutional neural network for modeling sentence pairs. Transactions of the Association for Computational Linguistics, 4(1), 2016.

[56] Z.-H. You, K. C. Chan, and P. Hu. Predicting protein-protein interactions from primary protein sequences using a novel multi-scale local feature representation scheme and the random forest. PLoS One, 10(5):e0125811, 2015.

[57] Z.-H. You, Y.-K. Lei, L. Zhu, J. Xia, and B. Wang. Prediction of protein-protein interactions from amino acid sequences with ensemble extreme learning machines and principal component analysis. In BMC bioinformatics, volume 14, page S10, 2013.

[58] Z.-H. You, L. Zhu, C.-H. Zheng, H.-J. Yu, S.-P. Deng, and Z. Ji. Prediction of protein-protein interactions from amino acid sequences using a novel multi-scale continuous and discontinuous feature set. In BMC bioinformatics, volume 15, page S9, 2014.

[59] K. Yugandhar and M. M. Gromiha. Protein–protein binding affinity prediction from amino acid sequence. Bioinformatics, 30(24):3583–3589, 2014.

[60] S. Zhang, J. Zhou, H. Hu, H. Gong, L. Chen, C. Cheng, and J. Zeng. A deep learning framework for modeling structural features of rna-binding protein targets. Nucleic acids research, 44(4):e32–e32, 2015.

[61] S.-W. Zhang, L.-Y. Hao, and T.-H. Zhang. Prediction of protein–protein interaction with pairwise kernel support vector machine. International journal of molecular sciences, 15(2):3220–3233, 2014.

[62] Y. Zhang, W. Chan, and N. Jaitly. Very deep convolutional networks for end-to-end speech recognition. In IEEE International Conference on Acoustics, Speech and Signal Processing (ICASSP), pages 4845–4849. IEEE, 2017.

[63] T. Zhou, M. Chen, J. Yu, and D. Terzopoulos. Attention-based natural language person retrieval. In IEEE Conference on Computer Vision and Pattern Recognition, pages 27–34. IEEE, 2017.

[64] H. Zhu, F. S. Domingues, I. Sommer, and T. Lengauer. Noxclass: prediction of protein-protein interaction types. BMC bioinformatics, 7(1):27, 2006.

[65] J. Zhu, H. Zou, S. Rosset, and T. Hastie. Multi-class adaboost. Statistics and its Interface, 2(3), 2009.

